# Differential roles of sleep spindles and sleep slow oscillations in memory consolidation

**DOI:** 10.1101/153007

**Authors:** Yina Wei, Giri P Krishnan, Maxim Komarov, Maxim Bazhenov

## Abstract

Sleep plays an important role in consolidation of recent memories. However, the mechanisms of consolidation remain poorly understood. In this study, using a realistic computational model of the thalamocortical network, we demonstrated that sleep spindles (the hallmark of N2 stage sleep) and slow oscillations (the hallmark of N3 stage sleep) both facilitate spike sequence replay as necessary for consolidation. When multiple memories were trained, the local nature of spike sequence replay during spindles allowed replay of the memories independently, while during slow oscillations replay of the weak memory was competing to the strong memory replay. This led to the weak memory extinction unless when sleep spindles (N2 sleep) preceded slow oscillations (N3 sleep), as observed during natural sleep. Our study presents a mechanistic explanation for the role of sleep rhythms in memory consolidation and proposes a testable hypothesis how the natural structure of sleep stages provides an optimal environment to consolidate memories.

**Significant Statement:** Numerous studies suggest importance of NREM sleep rhythms – spindles and slow oscillations - in sleep related memory consolidation. However, synaptic mechanisms behind the role of these rhythms in memory and learning are still unknown. Our new study predicts that sleep replay - the neuronal substrate of memory consolidation - is organized within the sleep spindles and coordinated by the Down to Up state transitions of the slow oscillation. For multiple competing memories, slow oscillations facilitated only strongest memory replay, while sleep spindles allowed a consolidation of the multiple competing memories independently. Our study predicts how the basic structure of the natural sleep stages provides an optimal environment for consolidation of multiple memories.

## Introduction

Sleep is believed to play an important role in consolidating of the recently learned memories (Walker and Stickgold, 2004; Diekelmann and Born, 2010; Born and Wilhelm, 2012; Rasch and Born, 2013). During sleep-related consolidation, memories become increasingly enhanced and resistant to interference(McGaugh, 2000). It was hypothesized that consolidation of memories during sleep occurs by reactivation of neuron ensembles engaged during learning. Indeed, spike sequence replay was observed in the neocortex(Euston et al., 2007; Ji and Wilson, 2007; Peyrache et al., 2009) following hippocampus-dependent tasks in coordination with replay in the hippocampus(Ji and Wilson, 2007) and following hippocampus-independent task (Ramanathan et al., 2015). Sequence replay during sleep is now believed to be a critical neural process involved in sleep dependent memory consolidation(Barnes and Wilson, 2014).

The natural sleep cycle consists of different sleep stages: Stage 1 (N1), Stage 2 (N2), Stage 3 (N3) of non-rapid eye movement (NREM) sleep, and rapid eye movement (REM) sleep(Rechtschaffen and Kales, 1968; Iber and Medicine, 2007; Silber et al., 2007). During NREM sleep, sleep spindles, 7-14 Hz brief bursts of rhythmic waves, are the most distinct oscillations that hallmark the N2 sleep(Loomis et al., 1935; Steriade et al., 1993a; Andrillon et al., 2011), while slow oscillations (<1 Hz), characterized by repetitive Up and Down states in all cortical neurons(Blake and Gerard, 1937; Steriade et al., 1993a; Steriade et al., 2001), are mainly observed during N3 sleep (also referred as slow wave sleep or deep sleep). While NREM sleep was shown to be particularly important for consolidating declarative (hippocampus-dependent) memories (Marshall et al., 2006; Mednick et al., 2013), human studies also suggest that NREM sleep may be involved in procedural (hippocampus-independent) memory consolidation, such as some simple motor tasks (Fogel and Smith, 2006). Indeed, selective depravation of N2 sleep, but not a REM sleep, led to the lower improvement for rotor pursuit task(Smith and MacNeill, 1994). Following a period of motor task learning, duration of NREM sleep(Fogel and Smith, 2006) and the number of sleep spindles (Morin et al., 2008) increased. The amount of performance increase in finger tapping task correlated with amount of NREM sleep (Walker et al., 2002), spindle density (Nishida and Walker, 2007) and delta power (Tamaki et al., 2013). Together studies suggests that NREM sleep is involved in consolidation of the simple motor tasks, while REM sleep may become critical for learning the more complex procedural memory tasks (see, e.g.: (Plihal and Born, 1997)). A recent animal study of consolidation of the hippocampus-independent (skilled upper-limb) memory reported that the reactivation of the neural activity was closely linked to the bursts of spindle activity and the waves of slow oscillation during NREM sleep. While it is evident that NREM sleep contributes to consolidation of memories through replay of spike sequences within neocortex, the mechanisms behind spike sequence replay and the functional role of characteristic NREM sleep rhythms – spindles and slow oscillation – in memory consolidation are still poorly understood.

Here we used a biophysically realistic model of the thalamocortical network, implementing synaptic plasticity (Wei et al., 2016) and effects of neuromodulators (Krishnan et al., 2016), to explore mechanisms of memory consolidation during NREM sleep. While we focus on the hippocampus-independent memory tasks, our results are also applicable to understanding the role of NREM sleep oscillations in consolidation of declarative memories. Our study revealed differential roles of sleep spindles and sleep slow oscillations in the spike sequence replay. We predict that sleep spindles and slow oscillations during NREM sleep play unique and complementary roles in consolidation of memories and that the natural sleep architecture, characterized by the well-defined sequence of sleep stages, is optimized to consolidate multiple mutually competing memories.

## Materials and Methods

### Model description

The thalamocortical network model incorporated thalamic relay (TC) and reticular (RE) neurons in the thalamus, as well as pyramidal neurons (PY) and inhibitory interneurons (IN) in the cortex(Bazhenov et al., 2002; Wei et al., 2016) organized with synaptic connectivity (Fig. 1). All neurons were modeled based on the Hodgkin-Huxley kinetics. The model included the changes in the level of the neuromodulators (reduction of acetylcholine and histamine; increase of GABA) to model transitions between awake state, N2 and N3 sleep states(Krishnan et al., 2016). The synaptic weights between cortical pyramidal cells were changed based on the spike timing-dependent synaptic plasticity (STDP) rule, as in (Wei et al., 2016). The units and description of parameters were summarized in Table 1.

**Figure 1.**
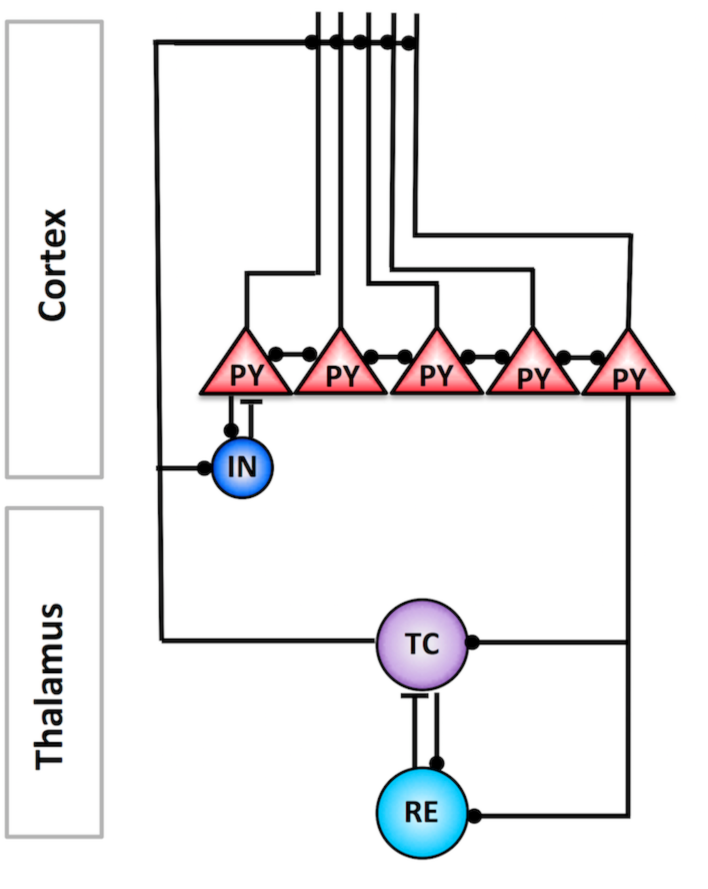
The schematic of the thalamocortical network model. The cortical layer was organized in a one-dimensional chain of pyramidal cells (PYs) and inhibitory neurons (INs). The thalamus model included thalamic relay (TC) and reticular thalamic (RE) neurons. Black filled circles and black bars represent excitatory and inhibitory connections between neurons, respectively.

**Table 1.** Main parameters. This table includes the units and description of the parameters used in the model.

| Parameters | Value | Description |
| --- | --- | --- |
| $C_m$ | $1 \mu\text{F}/\text{cm}^2$(TC;RE);<br>$0.75 \mu\text{F}/\text{cm}^2$(PY;IN) | Membrane capacitance |
| Thalamic cells |  |  |
| $S$ | $2.9 \times 10^{-4} \text{ cm}^2$ (TC); $1.43 \times 10^{-4} \text{ cm}^2$ (RE) | Area of neurons |
| $G_L$ | $0.01 \text{ mS}/\text{cm}^2$ (TC); $0.05 \text{ mS}/\text{cm}^2$ (RE) | Leakage conductance |
| $E_L$ | $-70 \text{ mV}$ (TC); $-77 \text{ mV}$ (RE) | Leakage reversal potential |
| $G_{KL}$ | $0.024 \text{ mS}/\text{cm}^2$ (TC); $0.012 \text{ mS}/\text{cm}^2$ (RE) | Potassium leakage conductance |
| $E_K$ | $-95 \text{ mV}$ (TC; RE) | Potassium reversal potential |
| $g_K$ | $10 \text{ mS}/\text{cm}^2$ (RE); $12 \text{ mS}/\text{cm}^2$ (TC) | Maximal potassium conductance |
| $g_{Na}$ | $90 \text{ mS}/\text{cm}^2$ (TC); $100 \text{ mS}/\text{cm}^2$ (RE) | Maximal sodium conductance |
| $g_T$ | $2.5 \text{ mS}/\text{cm}^2$ (TC); $2.2 \text{ mS}/\text{cm}^2$ (RE) | Low-threshold $\text{Ca}^{2+}$ conductance |
| $g_h$ | $0.016 \text{ mS}/\text{cm}^2$ (TC); $0 \text{ mS}/\text{cm}^2$ (RE); | Hyperpolarization-activated cation conductance |
| Cortical cells (Soma) |  |  |
| $S_{\text{soma}}$ | $1.0 \times 10^{-6} \text{ cm}^2$ (PY;IN) | Area of the axosomatic compartment |
| $g_K$ | $200 \text{ mS}/\text{cm}^2$ (PY;IN) | Maximal potassium conductance |
| $g_{Na}$ | $3000 \text{ mS}/\text{cm}^2$ (PY); $2500 \text{ mS}/\text{cm}^2$ (IN) | Maximal sodium conductance |
| $g_{Na(p)}$ | $15 \text{ mS}/\text{cm}^2$ (PY); $0 \text{ mS}/\text{cm}^2$ (IN); | Maximal persistent sodium conductance |
| Cortical cells (Dendrite) |  |  |
| $\rho$ | $165$ (PY); $50$ (IN) | $S_{\text{dend}} = \rho S_{\text{soma}}$ |
| $G_L$ | $0.009 \text{ mS}/\text{cm}^2$ (PY); $0.009 \text{ mS}/\text{cm}^2$ (IN) | Leakage conductance |
| $E_L$ | $-67 \text{ mV}$ (PY); $-70 \text{ mV}$(IN) | Leakage reversal potential |
| $G_{KL}$ | $0.011 \text{ mS}/\text{cm}^2$ (PY); $0.009 \text{ mS}/\text{cm}^2$ (IN) | Potassium leakage conductance |
| $E_K$ | $-95 \text{ mV}$ (PY;IN) | Potassium reversal potential |
| $g_{Na}$ | $0.8 \text{ mS}/\text{cm}^2$ (PY;IN) | Maximal sodium conductance |
| $g_{Na(p)}$ | $2.5 \text{ mS}/\text{cm}^2$ (PY); $0 \text{ mS}/\text{cm}^2$ (IN); | Maximal persistent sodium conductance |
| $g_{HVA}$ | $0.01 \text{ mS}/\text{cm}^2$ (PY;IN) | Maximal high-threshold $\text{Ca}^{2+}$ conductance |
| $g_{KCa}$ | $0.05 \text{ mS}/\text{cm}^2$ (PY;IN) | Slow $\text{Ca}^{2+}$-dependent $\text{K}^+$ conductance |
| $g_{Km}$ | $0.02 \text{ mS}/\text{cm}^2$ (PY); $0.015 \text{ mS}/\text{cm}^2$ (IN) | Slow voltage-dependent noninactivating $\text{K}^+$ conductance |

#### Neuromodulators and sleep stages

Our model implemented the change of neuromodulators, such as acetylcholine (ACh), histamine (HA), and GABA, in the intrinsic and synaptic currents to model transitions between sleep stages(Krishnan et al., 2016). Specifically, ACh modulates the potassium leak currents in all neurons and the strength of excitatory AMPA connections in the cortex; HA modulates the strength of the hyperpolarization-activated cation current (Ih) in TC neurons; GABA modulates the maximal conductance of the GABAergic synapses in IN and RE neurons. In the model, the reduction of ACh was implemented as an increase of potassium leak conductance in TC, PY and IN neurons, a reduction of potassium leak conductance in RE cells(McCormick, 1992), and an increase in AMPA connection strength(Kimura et al., 1999). The higher value of HA was implemented as a positive shift in the activation curve of a hyperpolarization-activated cation current (I_h_) (McCormick and Williamson, 1991; McCormick, 1992).

#### Intrinsic currents

The cortical PY and IN cells included dendritic and axo-somatic compartments, similar as previous studies(Timofeev et al., 2000; Bazhenov et al., 2002; Chen et al., 2012; Wei et al., 2016) and were described by Hodgkin-Huxley kinetics:
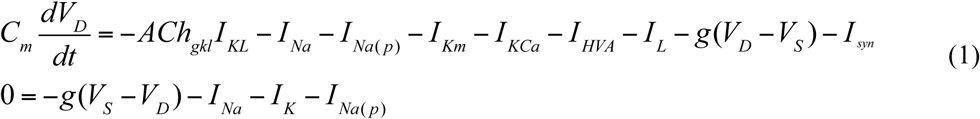

where ACh_gkl_ is inversely proportional to the level of ACh which modulates the potassium leak current (ACh_gkl_=0.133, 0.228 and 0.38 for awake, N2 and N3 sleep, respectively), *I_KL_* is a potassium leak current, *I_Na_* is a fast sodium current, *I_Na(P)_* is a persistent sodium current, *I_Km_* is a slow voltage-dependent non-inactivating potassium current, *I_KCa_* is a slow Ca^2+^-dependent K^+^ current, *I_HVA_* is a high-threshold Ca^2+^ current, I_L_ is the Cl^-^ leak current, g is the conductance between axo-somatic and dendritic compartment. *V_D_* and *V_S_* are the membrane potentials of dendritic and axosomatic compartments, and Isyn is the sum of synaptic currents to the neuron. The persistent sodium current *I_Na(P)_* was included in the axosomatic and dendritic compartment of PY cells to increase bursting propensity. IN cells had the same intrinsic currents as those in PY except that *I_Na(P)_* was not included. The ratio of dendritic area to axosomatic area was *ρ* =165 for PY, and *ρ* = 50 for IN. All the voltage-dependent ionic currents were of the form

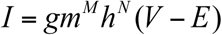

where g is the conductance, m and h are gating variables, V is the voltage of the corresponding compartment and E is the reversal potential. The detailed description of individual currents was provided in our previous study(Wei et al., 2016).

The thalamic TC and RE cells were modeled as a single compartment

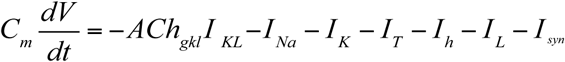

where ACh_gkl_ in TC cells is 0.4, 0.96, and 1.6 for awake, N2 and N3 sleep. ACh_gkl_ in RE cells is 0.9, 0.81, and 0.45 for awake, N2 and N3 sleep. *I_KL_* is a potassium leak current, *I_Na_* is a fast sodium current, *I_K_* is a fast potassium current, *I_T_* is a low threshold Ca^2+^ current, I_h_ is a hyperpolarization-activated cation current, IL is a Cl^-^ leak current, and Isyn is the sum of the synaptic currents to the neuron. The hyperpolarization-activated cation current *I_h_* was only included in TC neurons, not in RE neurons. The detailed description of individual currents was provided in our previous study(Wei et al., 2016). The effect of HA on I_h_ was implemented as a shift of HA_gh_ in the activation curve:

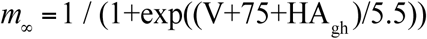

where HA_gh_ is -24 mV, -2 mV, -1mV for awake, N2 and N3 sleep, respectively.

#### Synaptic currents

The equations for GABA_A_, AMPA, and NMDA synaptic currents were described by first-order activation schemes, and the GABA_B_ synaptic currents had a more complex scheme of activation that involved the activation of K^+^ channels by G proteins(Destexhe et al., 1996). The equations for all synaptic currents used in this model were given in our previous studies(Bazhenov et al., 2002; Wei et al., 2016). In this paper, we added the level of ACh and GABA to modulate AMPA, and GABA_A_ synaptic currents as described by

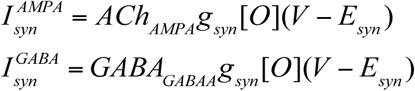

where g_syn_ is the maximal conductance, [O] is the fraction of open channels, and E_syn_ is the reversal potential (E_AMPA_=0 mV, E_NMDA_=0 mV, and E_GABAA_=-70 mV). ACh_AMPA_ is the variable that modulates AMPA synaptic currents for cortical PY-PY, TC-PY, and TC-IN connections by the level of ACh. ACh_AMPA_ from PY cells is 0.133, 0.1938, and 0.4332 for awake, N2 and N3 sleep. ACh_AMPA_ from TC cells is 0.6, 0.72 and 1.2 for awake, N2 and N3 sleep. GABA_GABAA_ is the variable that modulates GABA synaptic currents for cortical IN-PY, RE-RE and RE-TC connections. GABA_GABAA_ from IN cells is 0.22, 0.264 and 0.44 for awake, N2 and N3 sleep. GABA_GABAA_ from RE cells is 0.6, 0.72 and 1.2 for awake, N2 and N3 sleep, respectively.

The maximal conductance for each synapse were g_GABAA(RE-TC)_ = 0.06 μS, g_GABAB(RE-TC)_= 0.0025 μS, g_GABAA(RE-RE)_ = 0.1 μS, g_AMPA(TC-RE)_ = 0.06 μS, g_AMPA(TC-PY)_ = 0.14 μS, g_AMPA(TC-IN)_ = 0.12 μS, g_AMPA(PY-PY)_ = 0.24 μS, g_NMDA(PY-PY)_ = 0.01 μS, g_AMPA(PY-IN)_ = 0.12 μS, g_NMDA(PY-IN)_ = 0.01 μS., g_AMPA(PY-TC)_ = 0.04 μS, g_AMPA(PY-RE)_ = 0.08 μS and g_GABAA(IN-PY)_ = 0.24 μS.

In addition, spontaneous miniature EPSPs and IPSPs were implemented for PY-PY, PY-IN and IN-PY connections. The arrival times of spontaneous miniature EPSPs and IPSPs were modeled by Poisson processes(Stevens, 1993), with time-dependent mean rate μ = (2/(1+exp(-(t-t_0_)/F))-1)/250(Bazhenov et al., 2002), where t_0_ is a time instant of the last presynaptic spike(Timofeev et al., 2000). The mEPSP frequency (F) and amplitude (A) were F_PY-PY_ = 30, F_PY-IN_ = 30, F_IN-PY_ = 30, A_PY-PY_ =0.2 μS, A_PY-IN_=0.2 μS, and A_IN-PY_=0.2 μS.

#### Network geometry

Network architecture of the thalamocortical model was based on the previous studies of the spindle and slow oscillations(Bazhenov et al., 2002; Bonjean et al., 2011; Chen et al., 2012; Krishnan et al., 2016; Wei et al., 2016). The thalamocortical network consisted of five hundred cortical pyramidal (PY) neurons, one hundred cortical inhibitory (IN) neurons, one hundred thalamic relay (TC) neurons and one hundred thalamic reticular (RE) neurons. Each layer was organized in a one-dimensional chain, and the connectivity was local and determined by the radii of connections. The PY and IN neurons received AMPA and NMDA synapses from other PY neurons, and PY neurons also received GABA_A_ from IN neurons.

The radii of connections between cortical neurons were R_AMPA(PY-PY)_ = 5, R_NMDA(PY-PY)_ = 5, R_AMPA(PY-IN)_ = 1, R_NMDA(PY-IN)_ = 1 and R_GABAA(IN-PY)_= 5. The TC neurons projected to RE neurons through AMPA synapses (R_AMPA(TC-RE)_ =8), and connections from RE to TC neurons included GABA_A_ and GABA_B_ synapses (R_GABAA(RE-TC)_= 8, R_GABAB(RE-TC)_= 8). The radii of connections between RE and RE were R_GABAA(RE-RE)_ = 5. Thalamocortical connections were wider and mediated by AMPA synapses from TC neurons (R_AMPA(TC-PY)_ =15, _RAMPA(TC-IN)_ =3); corticothalamic connections were mediated AMPA synapses from PY neurons (R_AMPA(PY-TC)_ = 10, R_AMPA(PY-RE)_ =8).

### Statistical Analysis

The two-sample t-test was used for statistical analysis of differences between means from two samples. The paired t-test was used to compare the mean of normalized spatial correlation at different distance in spindles and slow oscillations. One-way ANOVA was used for comparisons across the multiple groups, with Bonferroni’s post hoc test was applied to determine the significant group. Two-way ANOVA was used for comparisons across groups with two independent factors (e.g. one factor is different training time of Seq2, and the second factor is Seq2 with or without the imprint of Seq1 in the network).

### Data Analysis

#### Sequence learning analysis

To model sequence learning, the model was presented with multiple trials of sequential input to the groups of selected cortical neurons. The performance of sequence recall was measured by the percentage of success of sequence recall during test sessions when only the first group of a sequence was stimulated. We applied a String Match (SM) method to measure the similarity between one sequence and an ideal sequence (e.g. “ABCDE”). The SM was calculated by 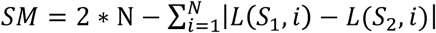, where *S*_1_ is the one sequence, *S*_2_ is the subset of ideal sequence that only contains the same elements as *S*_1_, *N* is the sequence length, *L*(*S*, *i*) represents the location of element i in a sequence S. SM was then normalized by dividing M, where M is the maximum SM calculated by two identical ideal sequences. The sequence recall was defined as successful when the SM between a recalled sequence and an ideal sequence was larger than 0.8, indicating 80% similarity.

#### Sequence replay measurement during sleep

The replay measure was calculated similar to the previous case 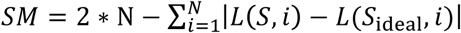, where *S*_ideal_ stands for correct sequence. The sequence S was defined with the following steps: (i) for each spike inside the first group *(e. g., n* ∊ [200,204]) we identified the neighboring (closest in time) spikes 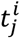 in all other neurons (i stands for group index, j stands for neuron index); (ii) for each group, we identified average firing times as 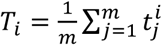; (iii) the sequence S was formed according to the order of average firing times *T_i_*. We rejected the sequences, whose total duration exceeded 100 ms.

#### Analysis of synaptic weights

Synaptic weights between neurons in a direction of sequence activation were enhanced due to the sequence replay. The mean of the changes of synaptic weights associated with a given sequence were used to characterize memory strength. Probability of enhancing Seq2 in two-sequence learning was calculated by counting the relative (over the total number of trials) number of trials that had a trend of increasing the mean synaptic weights associated with Seq2 for the last 100s of N3 sleep.

#### Spatiotemporal cluster analysis

Different contiguous activation regions were labeled based on connected components of smoothed spatio-temporal pattern (using **bwlabel** function in MATLAB). The region with the same-labeled number was considered as one cluster. The histogram of neuron number within each cluster was plotted during spindle and slow oscillations.

#### Computational methods

All model simulations were performed using a forth-order Runge-Kutta integration method with a time step of 0.02 ms. Source C++ was compiled on a Linux server using the G++ compiler. Part of the simulation was run on the Neuroscience Gateway(Sivagnanam et al., 2013). All data processing was done with custom-written programs in Matlab (MathWorks, Natick, MA).

## Results

### Spike sequence replay and synaptic mechanisms of memory consolidation during NREM sleep

We tested the role and the mechanisms of spike sequence replay for memory consolidation in a thalamocortical network model implementing awake, N2 and N3 sleep stages due to the variations in neuromodulators (Krishnan et al., 2016). The network model included thalamic relay (TC) and reticular (RE) neurons in the thalamus, as well as pyramidal neurons (PY) and inhibitory interneurons (IN) in the cortex(Fig. 1). Synaptic connections between cortical neurons were plastic and controlled by STDP similar to our recent study (Wei et al., 2016). We first simulated a basic sequence of sleep stages, including periods of awake, N2, N3 sleep and a second awake period following sleep by changing the level of neuromodulators (Fig. 2a). Since our study is focused on understanding the role of the sleep rhythms observed during non-rapid eye movement (NREM) sleep – spindles and slow oscillations – in memory consolidation, we avoided modeling N1 sleep or rapid eye movement (REM) sleep. The awake state included one training session and three test sessions: before training, after training before sleep, and after sleep (Fig. 2a). In the model, each sleep stage had its own characteristic pattern of electrical activity as observed *in vivo* (Fig. 2b). The neuronal activity during awake stage (Fig. 2b, *left*) showed no specific spatiotemporal pattern and random fluctuations in the local field potentials (LFP), reflecting desynchronized cell firing. The N2 sleep (Fig. 2b, *middle*) was characterized by sleep spindle oscillations, consisted of 7-14 Hz brief bursts of rhythmic waves that lasted 0.5-3 seconds and recurred every 2-20 seconds(Loomis et al., 1935; Steriade et al., 1993a; Andrillon et al., 2011), while N3 sleep (Fig.2b, *right*) was dominated by slow oscillations (<1 Hz), characterized by repetitive Up and Down states in all cortical neurons(Blake and Gerard, 1937; Steriade et al., 1993a; Steriade et al., 2001). We want to note that while we observed in the model the “waning” spindle activity at the beginning of the Up states of slow waves in N3 (Bazhenov et al., 2002), the overall spatio-temporal structure of the network activity during N3 sleep was dominated by the traveling slow waves and was very different from that during N2 sleep.

**Figure 2.**
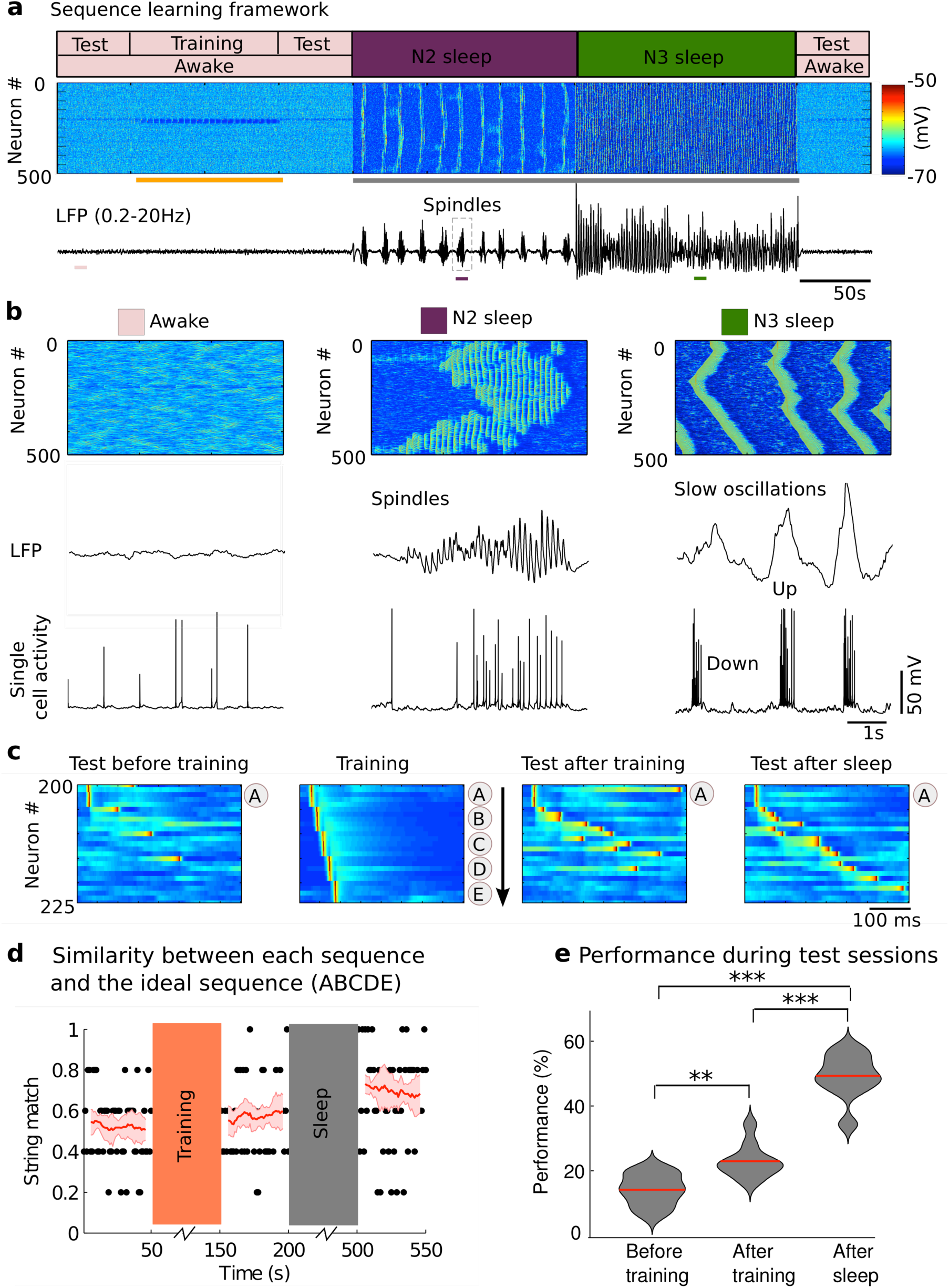
Network dynamics and sequence learning paradigm. **a)** The cortical network activity during transitions from awake state (pink block, *top*), to N2 sleep (purple block), to N3 sleep (dark green block) and back to awake. Raster plot (*middle*) shows membrane voltages of cortical pyramidal cells. Broadband filtered local field potential (LFP, *bottom*) from the cortical population. The sequence was learned during training period (orange bar). Grey bar represents the period of sleep. Performance was tested in three test sessions: before training, after training before sleep, and after sleep. **b**). The expanded view of characteristic spatio-temporal patterns (top), LFP (*middle*) and single cell activity of neuron #200 (*bottom*) during awake (*left*), N2 sleep (*middle*) and N3 sleep (*right*) from where pink, purple, dark green bars are shown in **a** (bottom). The spindle activity during N2 sleep revealed a typical waxing-waning pattern, consisted of 7-14 Hz brief bursts of rhythmic waves. The slow oscillations (<1Hz) during N3 sleep consisted of a typical Up and Down state transitions. **c**) The characteristic examples of a training session and three test sessions. The training included stimulating sequentially at groups A, B, C, D, and E. The test included stimulating only at group A (“pattern completion”). The sequence started at neuron #200. Each group included five neurons and it was stimulated for 10 ms. The delay between groups was 5 ms. **d**). Dot represent the string match between an ideal sequence (“ABCDE”) and each recalled sequence during test sessions for one trial. The value one represents a perfect match. The red line and the light red patch error bar represent mean and standard deviation of a moving average string match (window size=10) over all trials (n=10). **e**). The violin plot of the performance that was defined by the probability to recall a sequence (string match >=0.8) during multiple trials in each test session. Red line indicates mean value. Data were analyzed using one-way ANOVA with a Bonferroni post hoc test. * p<0.05, ** p<0.01, *** p<0.001.

During the training session, the model was presented with multiple trials of the input represented by a sequence of activation in selected groups of cortical neurons (Fig. 2c, *middle left*). Each group contained five neurons and was assigned a label (from A to E). By sequentially stimulating these five groups, the neuronal activity reflected sequential activation corresponding to the trained sequence, e.g., “ABCDE”. During test sessions (sequence recall), the model was only presented with the first input at group “A”. The characteristic examples of test sessions before training (Fig. 2c, *left*), after training before sleep (Fig.2c, *middle right*), and after sleep (Fig. 2c, *right*) showed a progressive increase of the correct sequence recall. To quantify memory recall performance, we used a string match (SM) measure (Fig. 2d, black dots), which measures the similarity between each recalled sequence during test sessions and the ideal sequence as trained, e.g. “ABCDE” (details in method section). We found that there was an overall increase of SM after training, and then after sleep (Fig. 2d, red line). We next calculated performance by measuring the success of a sequence recall – the percentage of correct sequence recalls (whose SM ≥ 0.8) when only first group of neurons (such as “A”) received stimulations (Fig. 2e). We observed a significant difference in the performance of sequence recall among all three test sessions as determined by one-way ANOVA (F_2,27_=103.19, p=2.26*10^−13^). The performance was significantly higher (p=0.0056, Bonferonni corrections) after repetitive training (22.6%±1.63%) compared to baseline performance before training (13.8%±1.53%), and was significantly higher after sleep (49.0%±2.18%) compared to before sleep (p=2.0174*10^−10^, Bonferonni corrections) (Fig. 2e).

To reveal network changes underlying performance increase, we next analyzed the dynamics of synaptic weights between cortical neurons. During the initial training phase, the ordered firing of neurons led to potentiation of synapses between neurons in the order of the trained sequence, while the synapses corresponding to the opposite order of the learned sequence were depressed (Fig. 3a, left). Importantly, the change in synaptic connections (Fig. 3a, *left,* grey box) was observed during N2 (Fig. 3a, *middle*) and continued in subsequent N3 sleep (Fig. 3a, *right*). Overall, we found a progressive increase in synaptic weights that strengthened the trained sequence (Fig. 3b, *left*) during sleep; this led to significant enhancement of performance after sleep (Fig.3c, *left*).

**Figure 3.**
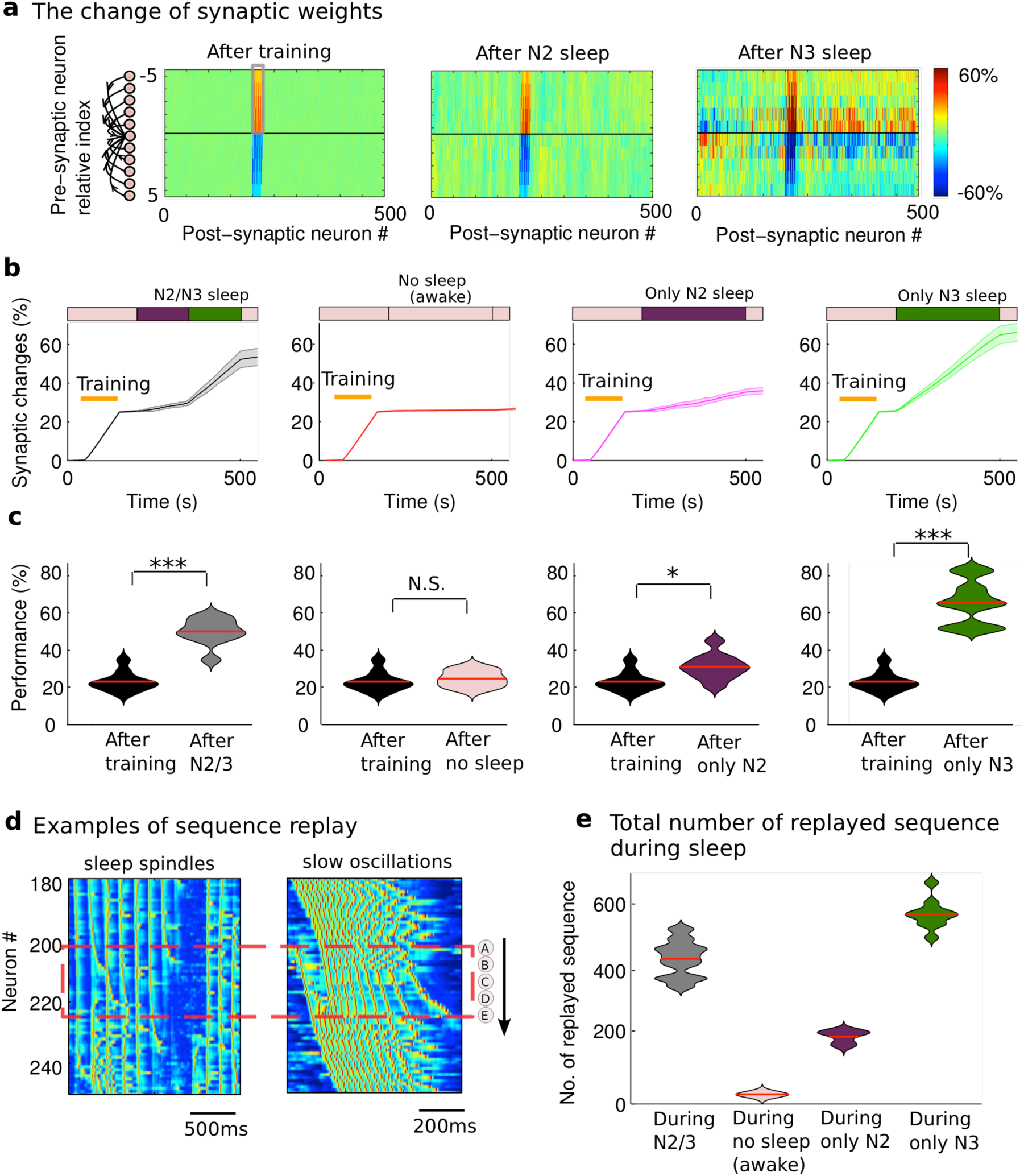
Spontaneous sequence replay mediates synaptic changes underlying memory consolidation during sleep. **a)** The change of synaptic weights relative to the initial values after training (*left*), N2 (*middle*) and N3 sleep (*right*). The synaptic weights between neurons in direction of sequence activation (grey box) were enhanced due to the sequence replay. **b**) The dynamics of the mean synaptic weights (grey box in **a**) shows the progressive increase in synaptic strength during normal N2/3 sleep (*left*), only N2 sleep (*middle right*); only N3 sleep (*right*). Note the lack of synaptic changes when sleep was supplemented by awake state of the same duration (*middle left*). Orange bar represents training period. The blocks in the top summarize the protocol of each experiment: Pink block - awake, purple block - N2 sleep, dark green block - N3 sleep. The patch error bar represents standard deviation. **c**) The violin plots of performance during test sessions after training (before sleep) and after sleep in four different experimental conditions corresponding to **b**. Red line indicates mean value. Data were analyzed using two-sample t test. * p<0.05, ** p<0.01, *** p<0.001. N.S. represents no significant difference. **d**) Characteristic examples of sequence (“ABCDE”) replay during sleep spindle and slow oscillations. **e**) The total count of replays (sequence “ABCDE”) during four difference experimental protocols.

In order to identify the role of different sleep stages in sequence learning, we next compared the change of synaptic weights and performance in four different conditions: 1) N2/N3 sleep (Fig. 3b and c, *left*); 2) No sleep (Fig. 3b and c, *middle left*); 3) only N2 sleep (Fig. 3b and c, *middle right*); 4) only N3 sleep (Fig. 3b and c, *right*). We found that recall performance of newly learned sequence was significantly enhanced after sleep (Fig. 3c) - either only N2 (t(9)=-2.9351, p = 0.0166, two-sample t-test), only N3 (t(9)=-11.7468, p = 9.2315*10^−7^, two-sample t-test), or N2/3 sleep (t(9)=-8.2644, p = 1.7056*10^−5^, two-sample t-test), but not after an equivalent awake period (t(9)=-0.6423, p = 0.5367, two-sample t-test; Fig. 3c, *middle left*). Synaptic potentiation in the direction of sequence learning represented a basic mechanism of performance improvement in all sleep conditions (Fig. 3b) and it was not significant during the equivalent awake period represented by the low level of background activity (Fig. 3b, *middle left*).

To reveal the neuronal mechanisms of synaptic reorganization during sleep, we analyzed the sequence reactivation during sleep of five groups of cortical neurons belonging to the sequence that was trained in awake (Fig. 3d, in the dotted red box). We found that the trained sequence was reactivated spontaneously during spindles (Fig. 3d, *left*) and slow oscillations (Fig. 3d, *right*). The total number of sequence replays during sleep was significantly higher compared to an equivalent awake period (Fig. 3e). We also observed a higher number of sequence replays during slow oscillation vs. spindles over the same period of sleep, which explains higher performance after N3 sleep alone vs. N2 sleep alone (compare Fig. 3c, middle/right and right). We concluded that spontaneous emergence of the sequence replay during sleep led to potentiation of synapses corresponding to the trained sequence and resulted in performance improvement after the sleep. Both sleep spindles and sleep slow oscillations provided the spike timing structure that was necessary for successful sequence replay and memory consolidation.

### Spatiotemporal nature of spindles and slow oscillations determines sequence replay

Is the sequence replay during sleep spindles different from that during slow oscillations? While both spindles and slow oscillations may activate neurons within the STDP time window to enable plastic changes, important difference seems to be in the overall spatio-temporal pattern. We first examined the cross-correlation of the Gaussian convoluted spike trains during spindles vs slow oscillations. When the peak of the cross-correlation was plotted for varying distances between network sites, its value reduced with increasing distance during both spindles and slow oscillations (Fig 4a-b). However, the amount of reduction during spindles (from 0.6 to 0.3) was much higher compared to slow oscillations (from 0.85 to 0.75). This difference was even more significant (Fig. 4c, black and red lines; t(149) =-116.1797, p = 4.5683*10^−148^, paired t-test) after normalization (Fig 4c) by the peak value of cross-correlation. This suggests that the spiking activities of the cortical neurons during spindles are correlated only within relatively small regions, while during slow oscillations activity across the network is globally coordinated due to its nature of the traveling wave propagation.

**Figure 4.**
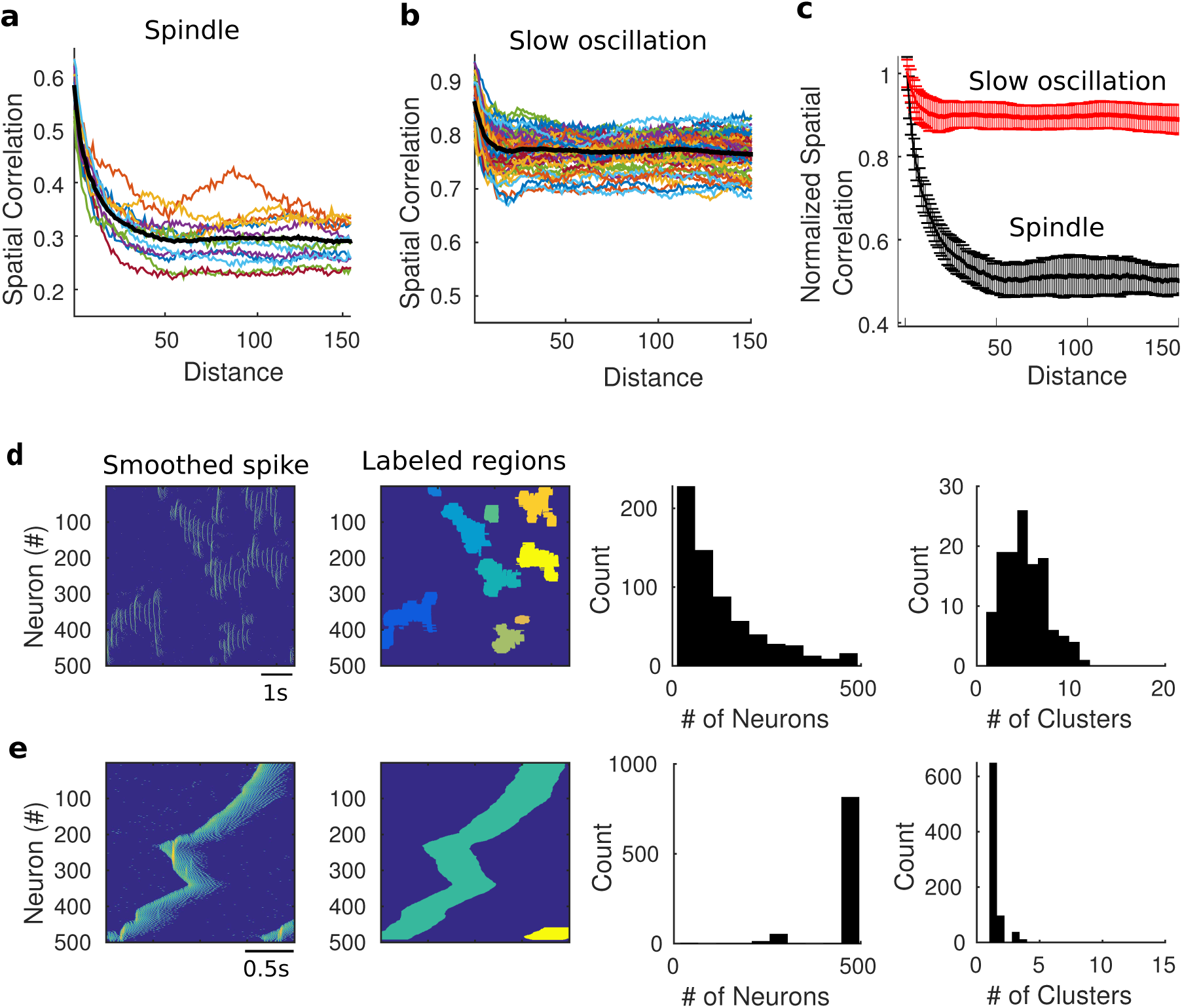
Differential spatiotemporal pattern of sleep spindles and slow oscillations. **a)** Spatial correlation between neurons at different distance during sleep spindles. **b**) Spatial correlation between neurons at different distance during slow oscillations. **c**) Normalized spatial correlation with mean and standard deviations. **d**,**e**) An example of smoothed spike trains and the clustered region, the histogram of neuron number that were identified within a cluster and the histogram of cluster numbers during spindle (**d**) and slow oscillations (**e**).

We further examined the local versus global nature of spindles and slow oscillations using a spatiotemporal cluster analysis. A typical single spindle event was built from many local clusters of neurons; while spiking was coordinated within each cluster, different clusters were semi-independent and initiated at and propagated from specific network location. In contrast, the slow waves had a more organized global structure with each wave initiated at one or just few locations and traveling over the entire network. To explore this difference, we applied clustering algorithm to count number of neurons within each cluster within a slow wave or a spindle event (Figs. 4d and e). While for slow oscillation a typical cluster included entire population of neurons (500 cells in our network), for sleep spindles a typical cluster size was much smaller. These results suggest that spindles may be better fitted to allow multiple memories to replay independently compare to the slow waves where multiple memories may have to interfere within one large cluster defined by the global pattern of a slow wave propagation. In the next we will show that this difference makes a large impact for the interplay of similar memories competing for the overlapping or closely located ensembles of neurons.

### The role of slow oscillation in organizing replay of multiple sequences

Brain can learn more than one motor sequences(Stephan et al., 2009). How do sleep rhythms coordinate multiple sequence replays? Here we show that spindles and slow oscillations compliment in their role in the consolidation of multiple sequences acquired in awake state. Specifically, we considered the case when the order of training of two sequences was opposite in the network topology and the neurons representing these sequences were relatively close in space, to explore the interaction between sequence replays during reactivation. Further, one sequence was trained longer (representing a strong memory) than another (representing a weak memory) to examine how two memories with different strength interact during sleep. The two sequences were learned by sequentially presenting stimuli at group A_1_, B_1_, C_1_, D_1_, E_1_ for Seq1, and at group E_2_, D_2_, C_2_, B_2_, A_2_ for Seq2, respectively (Fig. 5b). Seq1 “A_1_B_1_C_1_D_1_E_1_” was trained for 100s, representing a strong memory. Seq2 “E_2_D_2_C_2_B_2_A_2_” was trained for 40s, representing a weak memory. As before, recall performance for each sequence was measured based on the network responses by stimulating only first group of neurons in each sequence: group A_1_ (Fig. 5c), or group E_2_ (Fig. 5d).

**Figure 5.**
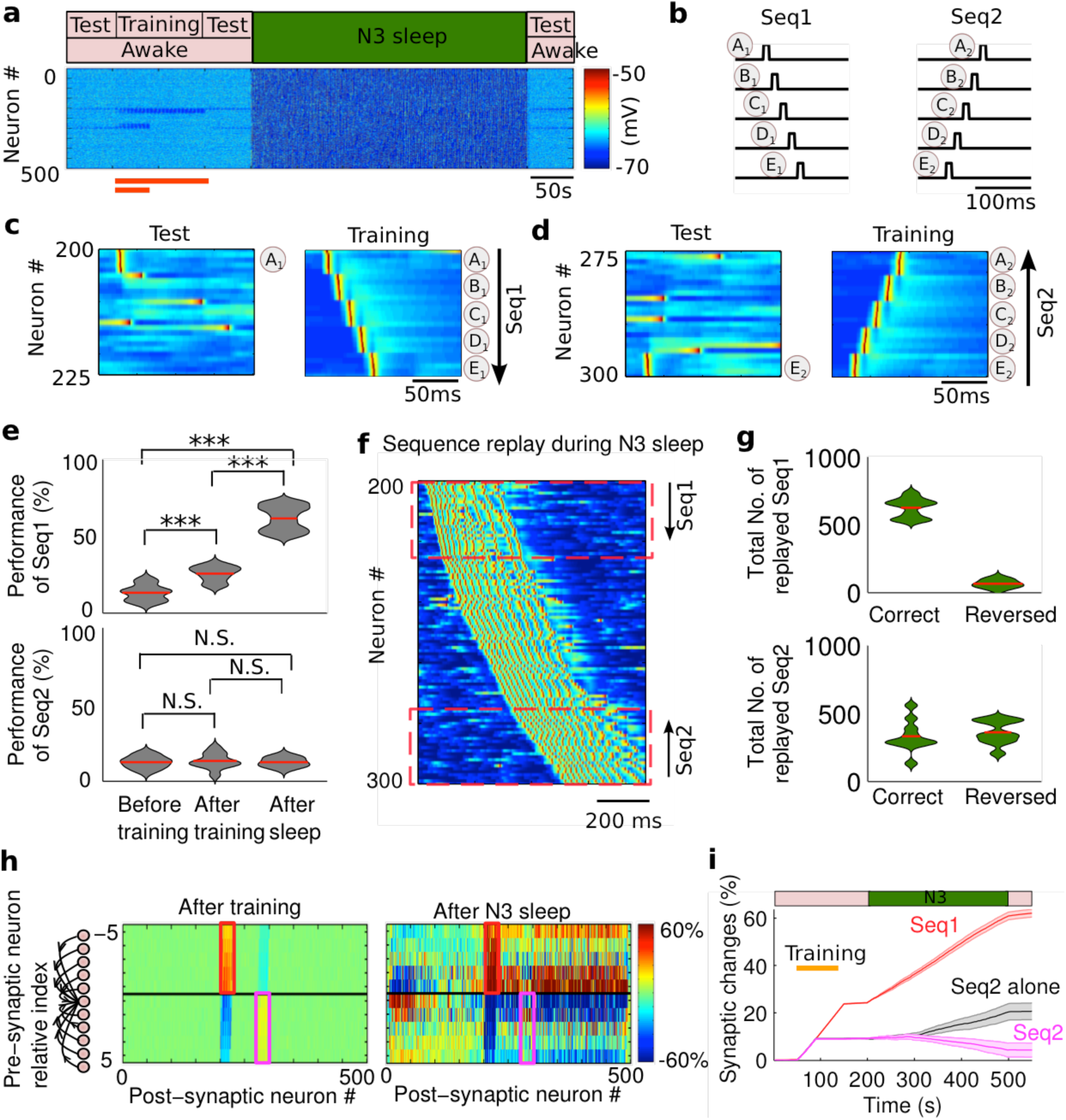
The role of slow oscillation during two-sequence learning. **a)** The model simulated transitions from awake to N3 sleep, and to awake again. Orange bar represents the duration of training of each sequence (*top:* Seq1; *bottom:* Seq2). **b**) A cartoon of the sequential network stimulation to generate two sequences. The duration of stimulation was 10ms for each group of neurons. The delay between subsequent stimuli of two groups was 5ms. Each group includes five neurons. **c**) Characteristic example of test and training of Seq1 (“A_1_B_1_C_1_D_1_E_1_”). The test was stimulating only at group A_1_. **d**) Test and training of Seq2 (“E_2_D_2_C_2_B_2_A_2_”). The test was stimulating only at group E_2_. The Seq1 and Seq2 started at neuron #200 and #300, respectively. **e**) The violin plot of performance for Seq1 and Seq2 during different test sessions. **f**) A characteristic example of the sequences replay during slow oscillations. **g**) The violin plot of the total replayed Seq1 (*top*) and Seq2 (*bottom*) during N3 sleep in correct and reverse order. The correct and reversed order for Seq1 were “A_1_B_1_C_1_D_1_E_1_” and “E_1_d_1_C_1_B_1_A_1_”, respectively. The correct and reversed order for Seq2 were “E_2_D_2_C_2_B_2_A_2_” and “A_2_B_2_C_2_D_2_E_2_”, respectively. **h)** The change of synaptic weights relative to the initial values after training (*left*) and after N3 sleep (*right*). Note that synaptic weights between neurons in the direction of Seq1 activation (red box) and Seq2 (magenta box) were both enhanced due to the training (*left*) but the effect decayed for Seq2 after N3 (*right*). **i)** The synaptic weights associated with Seq1 (red) were progressively increased during N3, while those associated with Seq2 (magenta) were decreased during N3 due to the interaction from the reactivation of Seq1. When Seq2 was trained alone (no interference) in the same experimental conditions, synaptic weights associated with Seq2 increased during N3 (black). The patch error bar represents standard error.

When only N3 sleep was simulated (Fig. 5a), we found a significant difference in the dynamics of the recall performance between strong (Seq1, F_2,27_ = 155.93, p=1.47*10^−15^, one-way ANOVA) and weak (Seq2, F_2,27_=0.13, p=0.8815, one-way ANOVA) memories. For Seq1, the performance was significantly increased (p=5.16*10^−4^, Bonferonni corrections) after training (25.6%±1.5720%) over the baseline (13.2%± 1.6653%), and further significantly improved (p=1.8014*10^−15^, Bonferonni corrections) after the sleep (61.6%±2.6297%). In contrast, the performance of the weakly trained Seq2 (Fig. 5e, *bottom*) was not significantly different from the baseline after initial training (13.2% ± 1.6111% vs. 12.4% ± 1.2579%, p=1, Bonferonni corrections) and was not significantly improved (p=1, Bonferonni corrections) after N3 sleep (12.4%±0.9333%). This reported change in performance was explained by the synaptic weight dynamics (Fig. 5h). During the initial training phase in awake, the ordered firing of neurons led to synaptic potentiation for the synapses associated with Seq1 (Fig. 5h, *left,* in the red box) and less significantly for the synapses associated with Seq2 (Fig. 5h, *left,* in the magenta box). During the following N3 sleep, synaptic connections associated with the strong memory were further increased (Fig. 5i, red line), however, those associated with the weak memory were reduced (Fig. 5i, magenta line). It is important to emphasize that in the absence of the strong memory (Seq1), the weak memory (Seq2) alone was enhanced (Fig. 5i, black line) during N3 sleep. These results are explained by the competition between two memories during slow oscillations: the strong memory was spontaneously reactivated in the correct order, while the weak memory was replayed more in the reversed order than in the correct one (Fig. 5f and g). The later was happening because of the global pattern of slow waves propagation controlled by the network sites associated with the strong memory. Therefore, slow oscillation preferentially enhanced the strong memory but impeded the consolidation of the weak memory.

To characterize how relative strength of the memory traces influences outcome of the sleep related consolidation, we varied the training duration of the Seq2 with or without the presence of Seq1 (a strong memory). For N3 sleep, synaptic changes associated with the Seq2 reversed the trend to decrease and even increased as the training duration of Seq2 increased (Fig. 6a). However, the amount of this increase was significantly less than when Seq2 was present alone (Fig. 6a, compare dark green and black lines; two-way ANOVA, F_1,126_=94.34, p=0). Recall performance of Seq2 after sleep was also significantly reduced in the presence of Seq 1 (Fig. 6b, compare dark green and black lines; two-way ANOVA, F_1,126_= 8.56, p=0.0041). Such difference was getting smaller as the strength of Seq2 increased (Fig. 6b). These results indicate that during slow oscillations, the presence of the strong memory trace, in close proximity to the cell population representing the weak memory, impede with the replay of the weak memory. Such interference was not observed when both memories were trained initially stronger (e.g., trained for 100sec or longer).

**Figure 6.**
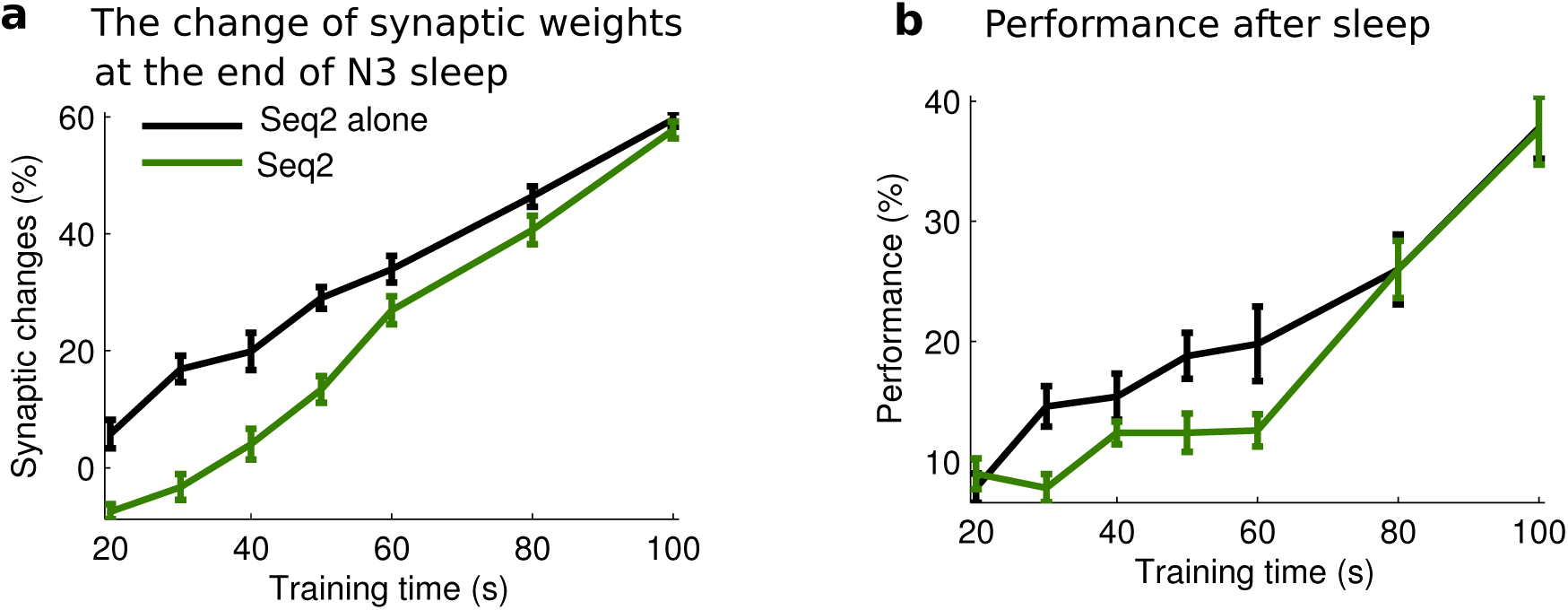
Effect of memory strength on the consolidation during slow oscillations. **a)** The dynamics of synaptic weights associated with Seq2 after N3 sleep for the different training duration (memory strength) of Seq2. Black line represents Seq2 trained alone. Dark green line indicates Seq2 trained along with the stronger Seq1. **b)** The change of Seq2 performance for the different training duration of Seq2. As the memory strength of Seq2 increased (longer training), the impact of interference on the synaptic weights and performance on Seq2 decreased.

### The role of sleep spindles in organizing replay of multiple sequences

We next tested the model with a sleep pattern similar to that of a natural sleep, where N3 was preceded by a period of N2 sleep (Fig. 7a). In this sleep conditions, the overall performance of both Seq1 (F_2,27_=527.81, p=2.28*10^−22^, one-way ANOVA) and Seq2 (F_2,27_=6.57, p=0.0047, oneway ANOVA) were enhanced following the sleep period. Post hoc analysis for Seq1 (Fig. 7b, *top*) indicated that performance was significantly increased (p=8.1761*10^−20^, Bonferonni corrections) after sleep (86%± 1.8379%) compared to that before sleep (25.6%± 1.5720%). For Seq2, the performance (Fig. 7b, *bottom*) also became significantly increased after sleep (27.6%±5.4062% vs. 13.2%± 1.6111%, p=0.01, Bonferonni corrections) due to the presence of 500s N2 sleep. We observed that both memories were reactivated more often in the correct order than in the reversed order during both N2 and N3 sleep (Fig. 7c and d). This led to a progressive increase in the synaptic weights associated with both sequences (Fig. 7e). The critical contribution of N2 sleep in memory consolidation was that during spindles synaptic weights representing correct order of firing increased both for the weak and strong memories (Fig. 7f). Such increase of synaptic weights associated with Seq2 during N2 sleep makes Seq2 stronger and resistant to further interference from the reactivation of Seq1 during the subsequent N3 sleep.

**Figure 7.**
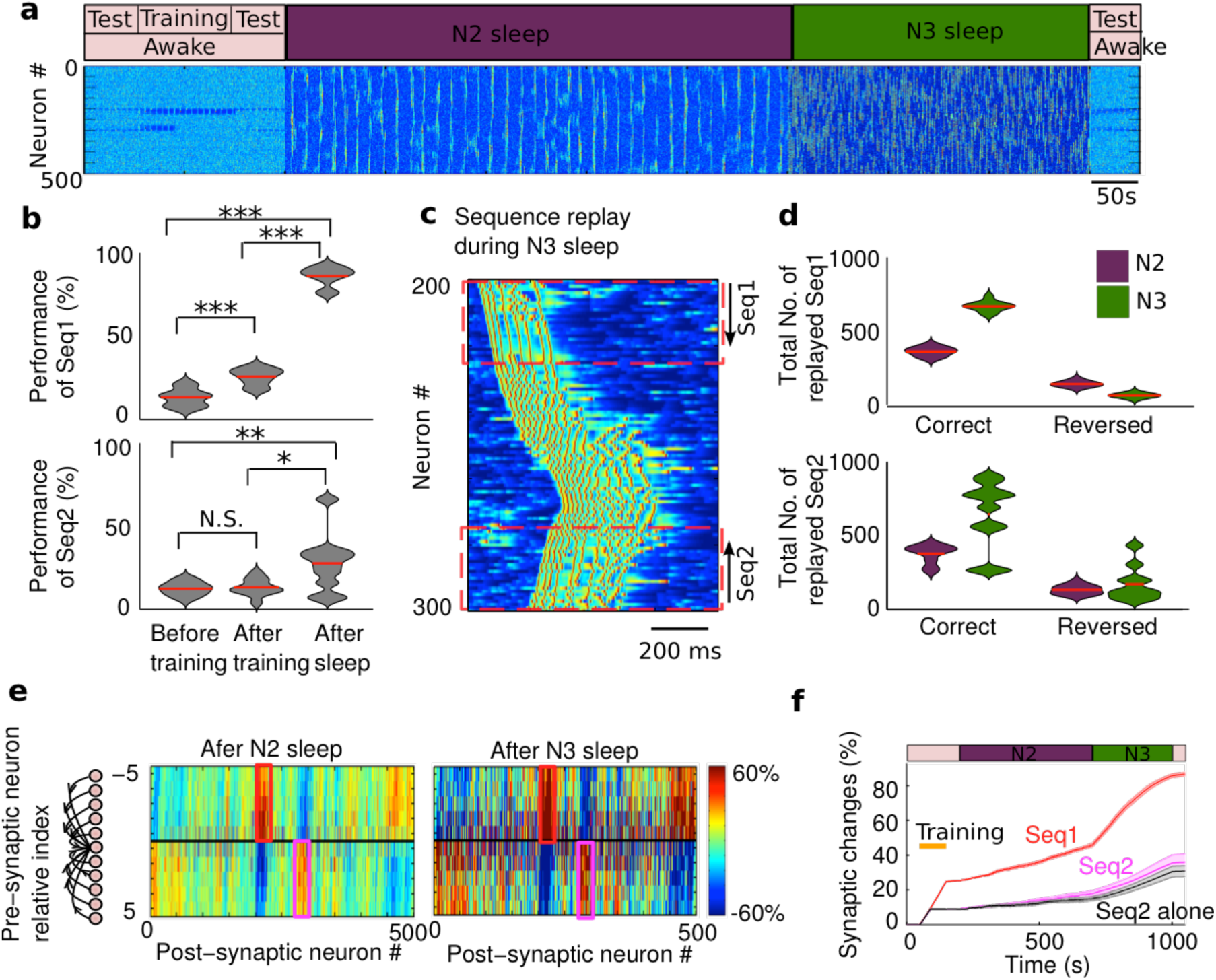
The role of sleep spindles during two-sequence learning. **a)** The model simulated transitions from awake to N2 sleep, to N3 sleep, and to awake again. Sequence training is the same as in Fig. 3. **b**) The violin plot of performance for Seq1 and Seq2 during test sessions. Note significant increase in Seq2 performance after the sleep. **c**) A characteristic example of sequence replay during slow oscillations. Note, that both Seq1 and Seq2 can be replayed during the same Up state of slow oscillation. **d**) The violin plot of the replay counts for Seq1 and Seq2 during N2 (purple) and N3 (dark green) sleep. Red line indicates mean value. Note that for both sequences number of correct order replays (“A_1_B_1_C_1_D_1_E_1_” for Seq1 and “E_2_D_2_C_2_B_2_A_2_” for Seq2) was higher than number of reversed order replays. Performance data were analyzed using one-way ANOVA with a Bonferroni post hoc test. * p<0.05, ** p<0.01, *** p<0.001. N.S. represents no significant difference. **e)** The change of synaptic weights relative to the initial values after N2 (*right*) and after subsequent N3 sleep (*left*). The synaptic change after training is the same as in **4j**). The enough amount of sleep spindles enhanced synaptic connections associated with both sequences independently. **f**). The progressive increase in synaptic weights associated with Seq1 (red), Seq2 (magenta), and Seq2 alone (black).

N2 sleep supported the consolidation of the weaker memory for the varying strength (training duration) of Seq2 (Fig. 8a, black and purple line, two-way ANOVA, F_1,126_=0.6, p=0.4393), except when the Seq2 was extremely weak and then sleep spindle activity was not able to mediate Seq2 replay. We found that given enough of the spindle activity early in the sleep cycle, synaptic weights associated with Seq2 enhanced sufficiently to allow for the further increase during the subsequent N3 sleep. Overall, there was no significant difference in the change of synaptic weights associated with Seq2 between two groups (with or without the presence of Seq1) after the full period of sleep (N2+N3) in this condition (Fig. 8b, two-way ANOVA, F_1,126_=0.84, p=0.3598). The performance of Seq2 recall after the full period of sleep also showed no significant difference whether Seq1 was present or Seq2 was trained alone (Fig. 8c, two-way ANOVA, F_1,126_=0.42, p=0.5192).

**Figure 8.**
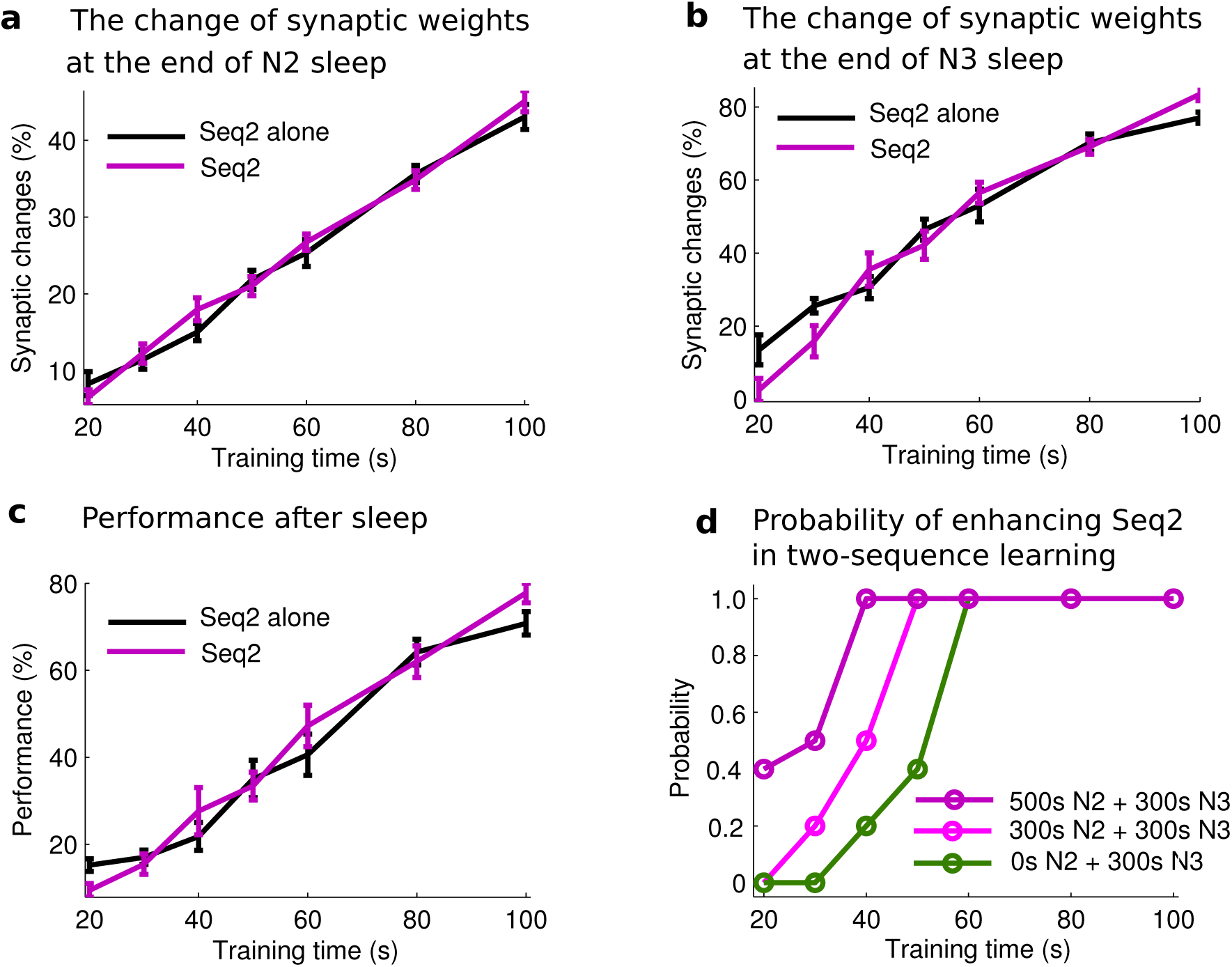
Effect of memory strength on the consolidation during normal N2/3 sleep. **a,b)**The change of synaptic weights associated with Seq2 after N2 (**a**) and N3 (**b**) sleep for the different training duration (memory strength) of Seq2. Importantly, after N2 sleep there is no difference in synaptic changes between Seq2 trained along with the stronger Seq1 and Seq2 trained alone. **c)** The change of Seq2 recall performance after N2/3 sleep for the different training duration of Seq2. **d**) Probability across trials of synaptic weights increase for Seq2, when trained along with Seq1, for different duration of N2 sleep.

We observed variability in the outcome of the Seq2 replay across individual trials, which was particularly high when Seq2 was weak (e.g., trained for 40s or less). Therefore, we next examined the probability (across trials) of improving performance of Seq2 in the presence of Seq1. Successful consolidation was defined as a trend of synaptic weights to increase during last 100 sec of sleep. As expected, increasing duration of initial Seq2 training, increased a probability of consolidation which saturated at 100% for experiments with training duration exceeding ~60 sec. Importantly, as the duration of N2 sleep increased, the probability of successful Seq2 consolidation also increased, shifting probability curves to the left (Fig. 8d). Thus, we conclude that sufficient amount of sleep spindles is necessary for the successful outcome of consolidation in experiments with a weak memory when other memories were imprinted in the same network (Fig. 8d).

## Discussion

In this study, using a realistic computational model of the thalamocortical network implementing sleep stages(Krishnan et al., 2016) and synaptic plasticity(Wei et al., 2016), we found that sleep spindles (7-14 Hz brief bursts of rhythmic waves) and sleep slow oscillations (<1 Hz rhythmic oscillations between Up and Down states of the thalamocortical network) both provide spatiotemporal pattern of cell firing that promotes spike sequence replay and organizes cortical spiking in a way optimal for synaptic plasticity and associated consolidation. Importantly, however, sleep spindles allowed independent and simultaneous replay of multiple memories, even when these memories were competing for the same or close ensembles of neurons. In contrast, sleep slow oscillation favored consolidation of the strong memories and could lead to the reverse replay and potentially to extinction of the weak memories. Taking into account that slow oscillation allowed faster rate of synaptic changes, our study predicts that a sequence of sleep stages N2 → N3, as observed during natural sleep in animals and humans, provides an optimal environment to reduce the interaction between memories during sleep replay and to maximize efficiency of consolidation.

### Mechanisms of spontaneous sequence replay

Synaptic plasticity is believed to be the cellular mechanism of learning and memory in the brain. Large body of studies support the idea that the spiking sequences of cortical neurons evoked by awake learning are replayed during sleep, leading to consolidation of memory (e.g., (Euston et al., 2007; Ji and Wilson, 2007; Peyrache et al., 2009; Ramanathan et al., 2015)). In our new study, neuronal sequences trained in awake, replayed spontaneously during sleep. In N2 sleep replay occurred during spindle events and was phase locked to the spindle oscillations, while in N3 it involved bursts of cortical cell firing during Up states of slow oscillations, consistent with the recent experimental findings(Ramanathan et al., 2015). We found no significant performance gain after an equivalent awake period, consistent with experimental studies(Fischer et al., 2002; Walker et al., 2002; Doyon et al., 2009; Tucker et al., 2016). We need to mention, however, that imposing a background synchronized activity, such as e.g., alpha rhythm in the quiet awake (Klimesch, 1999), could potentially lead to replay and consolidation. However, the study of consolidation in awake would go beyond the scope of this paper that is focused on the role of the major NREM sleep rhythms - sleep spindles and sleep slow oscillations - in memory consolidation.

Previous studies of the role of synaptic plasticity during sleep(Esser et al., 2007; Olcese et al., 2010) mainly focused on the global synaptic weights dynamics to investigate synaptic homeostasis (Tononi and Cirelli, 2003). In contrast, the focus of our study is local and task specific synaptic reorganization. Nevertheless, our study does not reveal global synaptic weights downscaling, as predicted by the synaptic homeostasis hypothesis (Tononi and Cirelli, 2003), and it is rather consistent with another line of evidences suggesting an increase of the learning specific synaptic weights during sleep (Timofeev and Chauvette, 2017).

Although we focused on the hippocampus-independent memory replay and thus our model does not include hippocampus, our results can be generalized to predict the role of NREM sleep in consolidation of the hippocampus-dependent memories. Hippocampal cell assembles reactivated during NREM sleep (Wilson and McNaughton, 1994; Skaggs and McNaughton, 1996) and spike sequence replay was reported to occur simultaneously in both hippocampus and cortex (Ji and Wilson, 2007) and it coincided with the hippocampal sharp-wave ripples (SWR)(Foster and Wilson, 2006; Wu and Foster, 2014). Hippocampal outflow during SWR coordinates reactivation of the relevant information distributed over multiple cortical modules(Schwindel and McNaughton, 2011). SWR tends to coincide with the transition from Down to Up states of slow oscillations(Battaglia et al., 2004) and cortical sequence replay (Peyrache et al., 2009), and may contribute to the initiation of the cortical Up states (Wei et al., 2016). Once initiated by the hippocampal input, replay in the cortical modules would be organized within patterns of sleep spindles and slow oscillation as predicted in our new study.

### Spindles and slow oscillations serve different roles in memory consolidation

Sleep spindles are a hallmark of N2 sleep, and shown to trigger neural plasticity and to contribute to memory consolidation by promoting synaptic short- and long-term potentiation(Rosanova and Ulrich, 2005). We found that sleep spindles alone were sufficient to facilitate synaptic changes in the model leading to performance improvement after the sleep. The performance gain was positively correlated with the amount of sleep spindles, in agreements with human studies (Walker et al., 2002; Fogel and Smith, 2006; Nishida and Walker, 2007; Morin et al., 2008; Astill et al., 2014; Laventure et al., 2016). Sleep slow oscillations are mainly observed during N3 sleep (deep sleep) and have been also associated with sleep-dependent performance enhancement. Enhancing slow oscillations by electrical stimulation improved the recall of word pairs in humans(Marshall et al., 2006). In our model, the period of sleep slow oscillations resulted in the improvement on the sequence completion task consistent with the previous experimental studies (Huber et al., 2004; Landsness et al., 2009; Astill et al., 2014).

One key difference, however, emerged between sleep spindles and slow oscillation on the nature of interaction between multiple memories. When network was trained for two close (in space) and opposite (directionally) sequences, spindle activity promoted the replay of both sequences independently or with little interaction, while slow oscillations led to the competitive interaction between sequence replays. The nature of this interaction depended on the properties of two trained sequences (similarity and orientation in space) and was especially prominent when one of the memories was weak (or trained for short time). In such case competition between two memories during slow oscillations (N3) could prevent the weak sequence from replay (and even force revered replay) and could lead to its extinction.

The model predicts that the difference between spatio-temporal patterns of sleep spindles vs sleep slow oscillation determined the role of spindles in minimizing interaction between similar memories during consolidation phase. The spindle activity was largely organized within small clusters of neurons. This allowed independent replay of many spike sequences simultaneously even when groups of neurons representing the sequences were close in space and were trained in the opposite direction. The slow oscillation was more widespread activity and showed a propagation pattern that may explain why it could lead to competition between sequences. For two sequences trained in the opposite network direction, the order of cell firing during slow waves frequently matched the order of the strong sequence but not the weak one. Only, when both sequences were sufficiently trained in awake state, they both mainly replayed in the correct order independently on the direction of slow wave traveling, in agreement with experimental data(Luczak et al., 2007).

This model prediction is consistent with many studies also reporting that sleep spindles emerge as local phenomena that are restricted to the specific brain regions involved in the recent task(Nishida and Walker, 2007; Tamaki et al., 2009; Andrillon et al., 2011; Nir et al., 2011). The spatio-temporal properties of the spindle activity may depend on the underlying cortical areas(Piantoni et al., 2017), with local and asynchronous spindles generated in deep cortical layers by the spatially restricted core thalamocortical system, while widespread spindles reflecting the distributed matrix system(Bonjean et al., 2012). Slow oscillation was shown to be a global travelling wave(Massimini et al., 2004). While coexistence of the active and silent cortical areas was reported during late sleep slow oscillation in some studies (Nir et al., 2011), this pattern was also found in our model, however it did not prevent competition.

Although spindles often co-occur with slow oscillations(Steriade et al., 1993b; Molle et al., 2011), our study is mainly focused on the differential role of sleep spindles vs slow oscillations in memory consolidation. We predict that for sleep spindles nested by slow waves during deep sleep the outcome of consolidation would be similar to what is reported here for slow oscillation. For mixed states including period of mainly spindle activity and some occasional slow waves, as commonly observed in humans during N2, the ratio of two will define the outcome.

During consolidation, new memory traces are stabilized or modified within the existing pool of memories(McGaugh, 2000; Diekelmann and Born, 2010). Memories may interfere with each other leading to forgetting (McKenzie and Eichenbaum, 2011; Robertson, 2012). Such interference has been observed between(Brashers-Krug et al., 1996; Brown and Robertson, 2007) and within memory domains (Baran et al., 2010; Brawn et al., 2013; Nettersheim et al., 2015). New learning was found to be particularly vulnerable to interference when competing learning events share stimulus features and when new events are trained in short temporal succession(Seitz et al., 2005; Yotsumoto et al., 2009). Interference may occur when one cluster of neurons “overwrites” or blocks the formation of another cluster of neurons. Sleep can protect memories from future interference(Ellenbogen et al., 2006), as well as rescue memories already damaged by interference(McDevitt et al., 2015a; McDevitt et al., 2015b). Our study predicts that sleep spindles may play a special role in protecting memories from interference and is also consistent with data of perceptual learning in humans(McDevitt et al., 2015b). We further predict that sleep spindles during N2 sleep and slow oscillations during N3 sleep play unique and complementary roles in consolidation of multiple memories and the order of stage 2 followed by stage 3 occurring during natural sleep is critical in preventing interference. Our study supports a hypothesis that the basic structure of sleep stages observed repeatedly across species from low vertebrates(Shein-Idelson et al., 2016) to humans(Rechtschaffen and Kales, 1968; Silber et al., 2007) provides an optimal environment for the consolidation of memories.

## CONFLICT OF INTEREST

None

## ACKNOWLEDGEMENTS

This work was supported by Office of Naval Research Multidisciplinary University Research Initiative Grant N000141612829 and National Institutes of Health Grants R01 EB009282 and R01 MH099645.

## Reference

Andrillon T, Nir Y, Staba RJ, Ferrarelli F, Cirelli C, Tononi G, Fried I (2011) Sleep Spindles in Humans: Insights from Intracranial EEG and Unit Recordings. The Journal of Neuroscience 31:17821.

Astill RG, Piantoni G, Raymann RJ, Vis JC, Coppens JE, Walker MP, Stickgold R, Van Der Werf YD, Van Someren EJ (2014) Sleep spindle and slow wave frequency reflect motor skill performance in primary school-age children. Front Hum Neurosci 8:910.

Baran B, Wilson J, Spencer RM (2010) REM-dependent repair of competitive memory suppression. Exp Brain Res 203:471–477.

Barnes DC, Wilson DA (2014) Slow-Wave Sleep-Imposed Replay Modulates Both Strength and Precision of Memory. The Journal of Neuroscience 34:5134–5142.

Battaglia FP, Sutherland GR, McNaughton BL (2004) Hippocampal sharp wave bursts coincide with neocortical “up-state” transitions. In: Learn Mem, pp 697–704.

Bazhenov M, Timofeev I, Steriade M, Sejnowski TJ (2002) Model of Thalamocortical Slow-Wave Sleep Oscillations and Transitions to Activated States. The Journal of Neuroscience 22:8691–8704.

Blake H, Gerard RW (1937) Brain potentials during sleep. American Journal of Physiology 119:692–703.

Bonjean M, Baker T, Lemieux M, Timofeev I, Sejnowski T, Bazhenov M (2011) Corticothalamic feedback controls sleep spindle duration in vivo. J Neurosci 31:9124–9134.

Bonjean M, Baker T, Bazhenov M, Cash S, Halgren E, Sejnowski T (2012) Interactions between core and matrix thalamocortical projections in human sleep spindle synchronization. J Neurosci 32:5250–5263.

Born J, Wilhelm I (2012) System consolidation of memory during sleep. Psychological Research 76:192–203.

Brashers-Krug T, Shadmehr R, Bizzi E (1996) Consolidation in human motor memory. Nature 382:252–255.

Brawn TP, Nusbaum HC, Margoliash D (2013) Sleep consolidation of interfering auditory memories in starlings. Psychol Sci 24:439–447.

Brown RM, Robertson EM (2007) Off-line processing: reciprocal interactions between declarative and procedural memories. J Neurosci 27:10468–10475.

Chen J-Y, Chauvette S, Skorheim S, Timofeev I, Bazhenov M (2012) Interneuron-mediated inhibition synchronizes neuronal activity during slow oscillation. The Journal of Physiology 590:3987–4010.

Cox R, Hofman WF, de Boer M, Talamini LM (2014) Local sleep spindle modulations in relation to specific memory cues. Neuroimage 99:103–110.

Destexhe A, Bal T, McCormick DA, Sejnowski TJ (1996) Ionic mechanisms underlying synchronized oscillations and propagating waves in a model of ferret thalamic slices. J Neurophysiol 76:2049–2070.

Diekelmann S, Born J (2010) The memory function of sleep. Nat Rev Neurosci 11:114–126.

Doyon J, Korman M, Morin A, Dostie V, Hadj Tahar A, Benali H, Karni A, Ungerleider LG, Carrier J (2009) Contribution of night and day sleep vs. simple passage of time to the consolidation of motor sequence and visuomotor adaptation learning. Exp Brain Res 195:15–26.

Ellenbogen JM, Hulbert JC, Stickgold R, Dinges DF, Thompson-Schill SL (2006) Interfering with Theories of Sleep and Memory: Sleep, Declarative Memory, and Associative Interference. Current Biology 16:1290–1294.

Esser SK, Hill SL, Tononi G (2007) Sleep Homeostasis and Cortical Synchronization: I. Modeling the Effects of Synaptic Strength on Sleep Slow Waves. Sleep 30:1617–1630.

Euston DR, Tatsuno M, McNaughton BL (2007) Fast-forward playback of recent memory sequences in prefrontal cortex during sleep. Science 318:1147–1150.

Fischer S, Hallschmid M, Elsner AL, Born J (2002) Sleep forms memory for finger skills. Proc Natl Acad Sci U S A 99:11987–11991.

Fogel SM, Smith CT (2006) Learning-dependent changes in sleep spindles and Stage 2 sleep. J Sleep Res 15:250–255.

Foster DJ, Wilson MA (2006) Reverse replay of behavioural sequences in hippocampal place cells during the awake state. Nature 440:680–683.

Huber R, Felice Ghilardi M, Massimini M, Tononi G (2004) Local sleep and learning. Nature 430:78–81.

Iber C, Medicine AAoS (2007) The AASM Manual for the Scoring of Sleep and Associated Events: Rules, Terminology and Technical Specifications: American Academy of Sleep Medicine.

Ji D, Wilson MA (2007) Coordinated memory replay in the visual cortex and hippocampus during sleep. Nat Neurosci 10:100–107.

Kimura F, Fukuda M, Tsumoto T (1999) Acetylcholine suppresses the spread of excitation in the visual cortex revealed by optical recording: possible differential effect depending on the source of input. Eur J Neurosci 11:3597–3609.

Klimesch W (1999) EEG alpha and theta oscillations reflect cognitive and memory performance: a review and analysis. Brain Res Brain Res Rev 29:169–195.

Krishnan GP, Chauvette S, Shamie I, Soltani S, Timofeev I, Cash SS, Halgren E, Bazhenov M (2016) Cellular and neurochemical basis of sleep stages in the thalamocortical network. eLife 5:e18607.

Landsness EC, Crupi D, Hulse BK, Peterson MJ, Huber R, Ansari H, Coen M, Cirelli C, Benca RM, Ghilardi MF, Tononi G (2009) Sleep-dependent improvement in visuomotor learning: a causal role for slow waves. Sleep 32:1273–1284.

Laventure S, Fogel S, Lungu O, Albouy G, Sévigny-Dupont P, Vien C, Sayour C, Carrier J, Benali H, Doyon J (2016) NREM2 and Sleep Spindles Are Instrumental to the Consolidation of Motor Sequence Memories. PLoS Biol 14:e1002429.

Loomis AL, Harvey EN, Hobart G (1935) Potential rhythms of the cerebral cortex during sleep. Science 81:597–598.

Luczak A, Bartho P, Marguet SL, Buzsaki G, Harris KD (2007) Sequential structure of neocortical spontaneous activity in vivo. Proc Natl Acad Sci U S A 104:347–352.

Marshall L, Helgadottir H, Molle M, Born J (2006) Boosting slow oscillations during sleep potentiates memory. Nature 444:610–613.

Massimini M, Huber R, Ferrarelli F, Hill S, Tononi G (2004) The sleep slow oscillation as a traveling wave. J Neurosci 24:6862–6870.

McCormick DA (1992) Neurotransmitter actions in the thalamus and cerebral cortex and their role in neuromodulation of thalamocortical activity. Prog Neurobiol 39:337–388.

McCormick DA, Williamson A (1991) Modulation of neuronal firing mode in cat and guinea pig LGNd by histamine: possible cellular mechanisms of histaminergic control of arousal. J Neurosci 11:3188–3199.

McDevitt E, Niknazar M, Mednick S (2015a) Sleep rescues perceptual learning from interference. Journal of Vision 15:1138–1138.

McDevitt EA, Duggan KA, Mednick SC (2015b) REM sleep rescues learning from interference. Neurobiol Learn Mem 122:51–62.

McGaugh JL (2000) Memory--a century of consolidation. Science 287:248–251.

McKenzie S, Eichenbaum H (2011) Consolidation and reconsolidation: two lives of memories? Neuron 71:224–233.

Mednick SC, McDevitt EA, Walsh JK, Wamsley E, Paulus M, Kanady JC, Drummond SPA (2013) The critical role of sleep spindles in hippocampal-dependent memory: a pharmacology study. J Neurosci 33:4494–4504.

Molle M, Bergmann TO, Marshall L, Born J (2011) Fast and slow spindles during the sleep slow oscillation: disparate coalescence and engagement in memory processing. Sleep 34:1411–1421.

Morin A, Doyon J, Dostie V, Barakat M, Hadj Tahar A, Korman M, Benali H, Karni A, Ungerleider LG, Carrier J (2008) Motor sequence learning increases sleep spindles and fast frequencies in post-training sleep. Sleep 31:1149–1156.

Nettersheim A, Hallschmid M, Born J, Diekelmann S (2015) The role of sleep in motor sequence consolidation: stabilization rather than enhancement. J Neurosci 35:6696–6702.

Nir Y, Staba RJ, Andrillon T, Vyazovskiy VV, Cirelli C, Fried I, Tononi G (2011) Regional Slow Waves and Spindles in Human Sleep. Neuron 70:153–169.

Nishida M, Walker MP (2007) Daytime naps, motor memory consolidation and regionally specific sleep spindles. PLoS One 2:e341.

Olcese U, Esser SK, Tononi G (2010) Sleep and Synaptic Renormalization: A Computational Study. In: J Neurophysiol, pp 3476–3493. Bethesda, MD.

Peyrache A, Khamassi M, Benchenane K, Wiener SI, Battaglia FP (2009) Replay of rule-learning related neural patterns in the prefrontal cortex during sleep. Nat Neurosci 12:919–926.

Piantoni G, Halgren E, Cash SS (2017) Spatiotemporal characteristics of sleep spindles depend on cortical location. Neuroimage 146:236–245.

Plihal W, Born J (1997) Effects of early and late nocturnal sleep on declarative and procedural memory. Journal of cognitive neuroscience 9:534–547.

Ramanathan DS, Gulati T, Ganguly K (2015) Sleep-Dependent Reactivation of Ensembles in Motor Cortex Promotes Skill Consolidation. PLOS Biology 13:e1002263.

Rasch B, Born J (2013) About sleep’s role in memory. Physiological reviews 93:681–766.

Rasch B, Büchel C, Gais S, Born J (2007) Odor Cues During Slow-Wave Sleep Prompt Declarative Memory Consolidation. Science 315:1426–1429.

Rechtschaffen A, Kales A (1968) A manual of standardized terminology, techniques and scoring system of sleep stages in human subjects. Bethesda, Md.: U.S. National Institute of Neurological Diseases and Blindness, Neurological Information Network.

Robertson EM (2012) New insights in human memory interference and consolidation. Curr Biol 22:R66–71.

Rosanova M, Ulrich D (2005) Pattern-specific associative long-term potentiation induced by a sleep spindle-related spike train. J Neurosci 25:9398–9405.

Rudoy JD, Voss JL, Westerberg CE, Paller KA (2009) Strengthening Individual Memories by Reactivating Them During Sleep. Science 326:1079–1079.

Schwindel CD, McNaughton BL (2011) Hippocampal-cortical interactions and the dynamics of memory trace reactivation. Prog Brain Res 193:163–177.

Seitz AR, Yamagishi N, Werner B, Goda N, Kawato M, Watanabe T (2005) Task-specific disruption of perceptual learning. Proceedings of the National Academy of Sciences of the United States of America 102:14895–14900.

Shein-Idelson M, Ondracek JM, Liaw HP, Reiter S, Laurent G (2016) Slow waves, sharp waves, ripples, and REM in sleeping dragons. Science 352:590–595.

Silber MH, Ancoli-Israel S, Bonnet MH, Chokroverty S, Grigg-Damberger MM, Hirshkowitz M, Kapen S, Keenan SA, Kryger MH, Penzel T, Pressman MR, Iber C (2007) The visual scoring of sleep in adults. J Clin Sleep Med 3:121–131.

Sivagnanam S, Majumdar A, Yoshimoto K, Astakhov V, Bandrowski A, Martone M, Carnevale NT (2013) Introducing the Neuroscience Gateway. In: Proceedings of the 5th International Workshop on Science Gateways. Zurich,Switzerland.

Skaggs WE, McNaughton BL (1996) Replay of neuronal firing sequences in rat hippocampus during sleep following spatial experience. Science 271:1870–1873.

Smith C, MacNeill C (1994) Impaired motor memory for a pursuit rotor task following Stage 2 sleep loss in college students. J Sleep Res 3:206–213.

Stephan MA, Meier B, Orosz A, Cattapan-Ludewig K, Kaelin-Lang A (2009) Interference during the implicit learning of two different motor sequences. Exp Brain Res 196:253–261.

Steriade M, Nunez A, Amzica F (1993a) Intracellular analysis of relations between the slow (< 1 Hz) neocortical oscillation and other sleep rhythms of the electroencephalogram. J Neurosci 13:3266–3283.

Steriade M, Timofeev I, Grenier F (2001) Natural Waking and Sleep States: A View From Inside Neocortical Neurons.

Steriade M, Contreras D, Curro Dossi R, Nunez A (1993b) The slow (< 1 Hz) oscillation in reticular thalamic and thalamocortical neurons: scenario of sleep rhythm generation in interacting thalamic and neocortical networks. J Neurosci 13:3284–3299.

Stevens CF (1993) Quantal release of neurotransmitter and long-term potentiation. Cell 72 Suppl:55–63.

Tamaki M, Matsuoka T, Nittono H, Hori T (2009) Activation of fast sleep spindles at the premotor cortex and parietal areas contributes to motor learning: A study using sLORETA. Clinical Neurophysiology 120:878–886.

Tamaki M, Huang T-R, Yotsumoto Y, Hämäläinen M, Lin F-H, Náñez JE, Watanabe T, Sasaki Y (2013) Enhanced spontaneous oscillations in the supplementary motor area are associated with sleep-dependent offline learning of finger-tapping motor-sequence task. Journal of Neuroscience 33:13894–13902.

Timofeev I, Chauvette S (2017) Sleep slow oscillation and plasticity. Curr Opin Neurobiol 44:116–126.

Timofeev I, Grenier F, Bazhenov M, Sejnowski TJ, Steriade M (2000) Origin of Slow Cortical Oscillations in Deafferented Cortical Slabs. Cerebral Cortex 10:1185–1199.

Tononi G, Cirelli C (2003) Sleep and synaptic homeostasis: a hypothesis. Brain Res Bull 62:143–150.

Tucker MA, Nguyen N, Stickgold R (2016) Experience Playing a Musical Instrument and Overnight Sleep Enhance Performance on a Sequential Typing Task. PLoS One 11:e0159608.

Walker MP, Stickgold R (2004) Sleep-dependent learning and memory consolidation. Neuron 44:121–133.

Walker MP, Brakefield T, Morgan A, Hobson JA, Stickgold R (2002) Practice with sleep makes perfect: sleep-dependent motor skill learning. Neuron 35:205–211.

Wei Y, Krishnan GP, Bazhenov M (2016) Synaptic Mechanisms of Memory Consolidation during Sleep Slow Oscillations. The Journal of Neuroscience 36:4231–4247.

Wilson MA, McNaughton BL (1994) Reactivation of hippocampal ensemble memories during sleep. Science 265:676–679.

Wu X, Foster DJ (2014) Hippocampal Replay Captures the Unique Topological Structure of a Novel Environment. In: J Neurosci, pp 6459–6469.

Yotsumoto Y, Chang LH, Watanabe T, Sasaki Y (2009) Interference and feature specificity in visual perceptual learning. Vision research 49:2611–2623.

